# Gender bias in research teams and the underrepresentation of women in science

**DOI:** 10.1101/741694

**Authors:** Roslyn Dakin, T. Brandt Ryder

**Affiliations:** Department of Biology, Carleton University, Ottawa, ON, K1S 5B6, Canada; Bird Conservancy of the Rockies, Fort Collins, Colorado, 80525, USA; Migratory Bird Center, Smithsonian Conservation Biology Institute, National Zoological Park, Washington, DC 20013, USA

**Keywords:** team science, collaboration, social learning, meta-science, scientific publishing, bibliometrics, higher education

## Abstract

Why are females still underrepresented in science? The social factors that affect career choices and trajectories are thought to be important but are poorly understood. We analyzed author gender in a sample of >61,000 scientific articles in the biological sciences to evaluate the factors that shape the formation of research teams. We find that authorship teams are more gender-assorted than expected by chance, with excess homotypic assortment accounting for up to 7% of published articles. One possible mechanism that could explain gender assortment and broader patterns of female representation is that women may focus on different research topics than men (i.e., the “topic preference” hypothesis). An alternative hypothesis is that researchers may consciously or unconsciously prefer to work within same-gender teams (the “gender homophily” hypothesis). Using network analysis, we find no evidence to support the topic preference hypothesis, because the topics of female-authored articles are no more similar to each other than expected within the broader research landscape. Instead, consistent with a model of moderate gender homophily, we find that the prevalence of matched-gender teams increases as a discipline moves towards gender parity. This can occur because latent preferences are more easily fulfilled in a gender-diverse environment. Finally, we show that female authors pay a substantial citation cost to work in gender-matched teams. Notably, the prevalence of homotypic assortment is predicted to increase in the future if more disciplines shift towards gender parity. These data indicate that social preferences can have important downstream consequences for the retention of women and other underrepresented groups in science.

---

The underrepresentation of women in science careers persists even in fields like biology where female undergraduate representation has exceeded parity (50%) for more than 30 years [1–3]. Some of the factors hypothesized to contribute to this leaky pipeline include gender differences in academic preferences and performance, the incompatibility of child-bearing with advanced training in the sciences, gender stereotypes, and discrimination [4–6]. Because science is increasingly a team endeavour, the social environment is expected to play a key role in career trajectories, collaboration opportunities, and subsequent retention [6–9]. However, analyses of collaboration and patterns of co-authorship, a key metric of academic success, are relatively scarce.

To analyze the gender composition of authorship teams, we compiled a sample of >101,000 scientific articles by multi-authored teams from the US, England, and Canada. This dataset represents 10% of the research output in those countries within seven major disciplines of biology (biophysics, ecology, plant sciences, biochemistry, neurosciences, cell biology and psychology; the sampled articles were from 2007, 2011, 2015 and 2018). Each of these disciplines also exhibits consistent differences in the overall representation of female authors (Fig. 1A). We used a large database of over 180 million birth records to match author first names with gender, yielding >61,000 articles for which both the first and last authors could be gender-identified with a high degree of certainty [10,11]. Our analysis focuses on these two author positions because most papers have at least two coauthors, and because the convention in the biological sciences is that the first and last authors made the greatest intellectual contribution to the research [10]. We find that the gender composition of authorship teams is not balanced: there are significantly more female-biased and male-biased author teams than expected by chance, based on the gender ratios within each discipline (Fig. 1B [12]).

**Figure 1.**
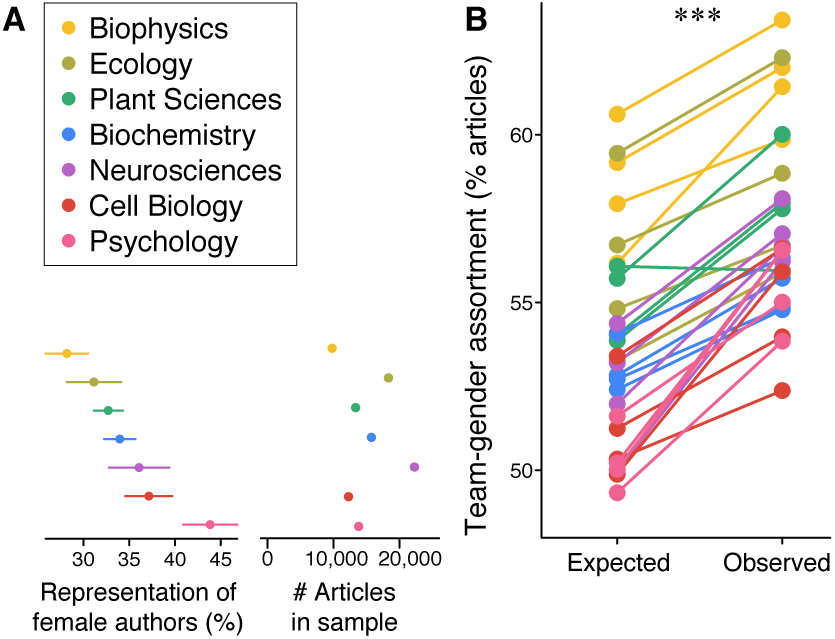
Scientific authorship teams are gender-assorted. A) We analyzed author gender across seven biological disciplines that vary in the representation of female authors (± SD). The intra-class correlation coefficient for representation was 0.78 [95% confidence interval = 0.50-0.95] indicating strong and consistent differences among disciplines (n = 28 discipline-years). B) The gender assortment of authorship teams was greater than expected (p < 0.0001). The null expectation is that team members would be chosen proportional to gender representation in the relevant discipline.

Why should authors work in gender-matched teams? One possible explanation is that females and males may prefer different research topics [1,7]. Hence, homotypic gender assortment may be a by-product of shared topic preferences. This “topic preference hypothesis” is intuitively appealing, because it could also explain the broader differences among scientific disciplines (Fig. 1A, [7]). We used a semantic network analysis to test this hypothesis, arranging the articles within each discipline into a network based on their shared keywords (Fig. 2A-B).

**Figure 2.**
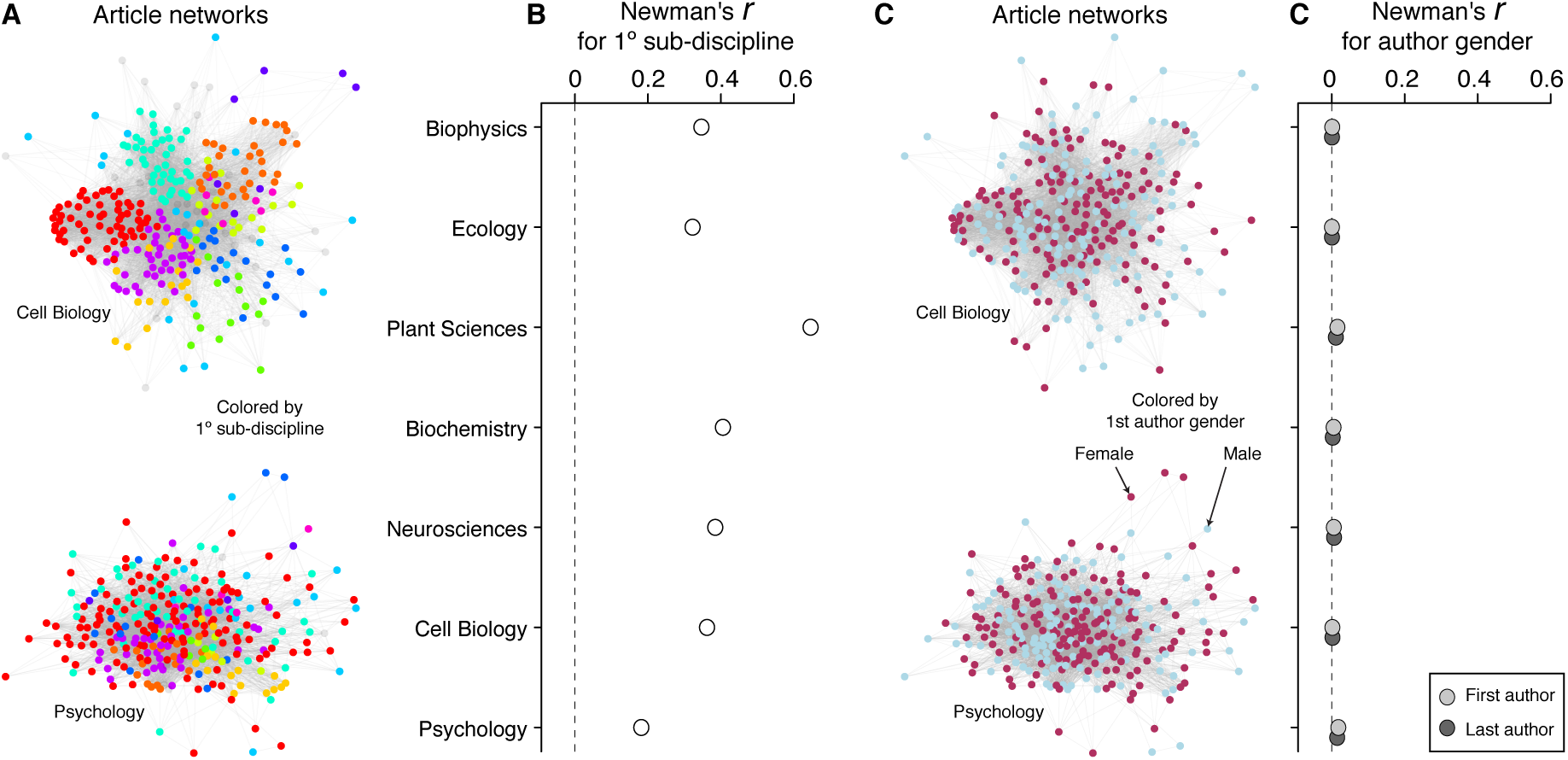
Gender is not associated with particular research topics within disciplines. A) We built an article network for each discipline containing all publications that could be linked by their shared keywords. Hence, each network represents a landscape of research topics. Two example networks are shown for Cell Biology and Psychology; each node represents an article, colored by its primary sub-discipline. B) We used Newman’s r to verify that the resulting networks were structured by sub-discipline. As expected, all of the networks had strong positive assortment (that is, the articles clustered near others in the same sub-discipline). C–D) We used these networks to test the hypothesis that male and female authors prefer different types of research. D) Contrary to this hypothesis, there was no clustering by author gender. All of the Newman’s r values for author gender were between 0 and 0.02, indicating that the female and male researchers were well-mixed with respect to research topics within each discipline. In B and D, the 95% confidence intervals were determined by bootstrapping, and the confidence intervals are much narrower than the data points. In A and C, only a random subset of 300 articles are shown for clarity. However, the analysis was based on the full networks (n = 4,102– 12,543 articles per network).

We then tested whether first (or last) author genders were clustered together within these networks, by examining network assortment of author gender (Fig. 2C-D). Contrary to the topic preference hypothesis, female-authored articles were not more semantically-related to each other than expected by chance (and nor were the male-authored articles; all r < 0.02). Moreover, we did not find any assortment by first-or last-author gender when examining networks that were pooled across the disciplines within each studied year (all r < 0.02).

An alternative explanation for homotypic gender assortment may be homophily, or a preference for same-gender coauthors [12–15]. Gender homophily is part of a broader mechanism of social selection, defined as when individuals form social relationships on the basis of preferred partner characteristics [16]. It is important to note that these preferences may often be unconscious (i.e., researchers need not be aware of the social preferences driving their career choices). To explore whether a model of homophily was consistent with the bibliometric data, we used an individual-based model to simulate a constant, two-fold greater preference for same-gender coauthors. The simulation was agnostic as to whether gender preferences were conscious. Remarkably, in both the simulation and the actual data, the gender assortment of teams increased markedly as female representation within a discipline approached parity (Fig. 3A-B).

**Figure 3.**
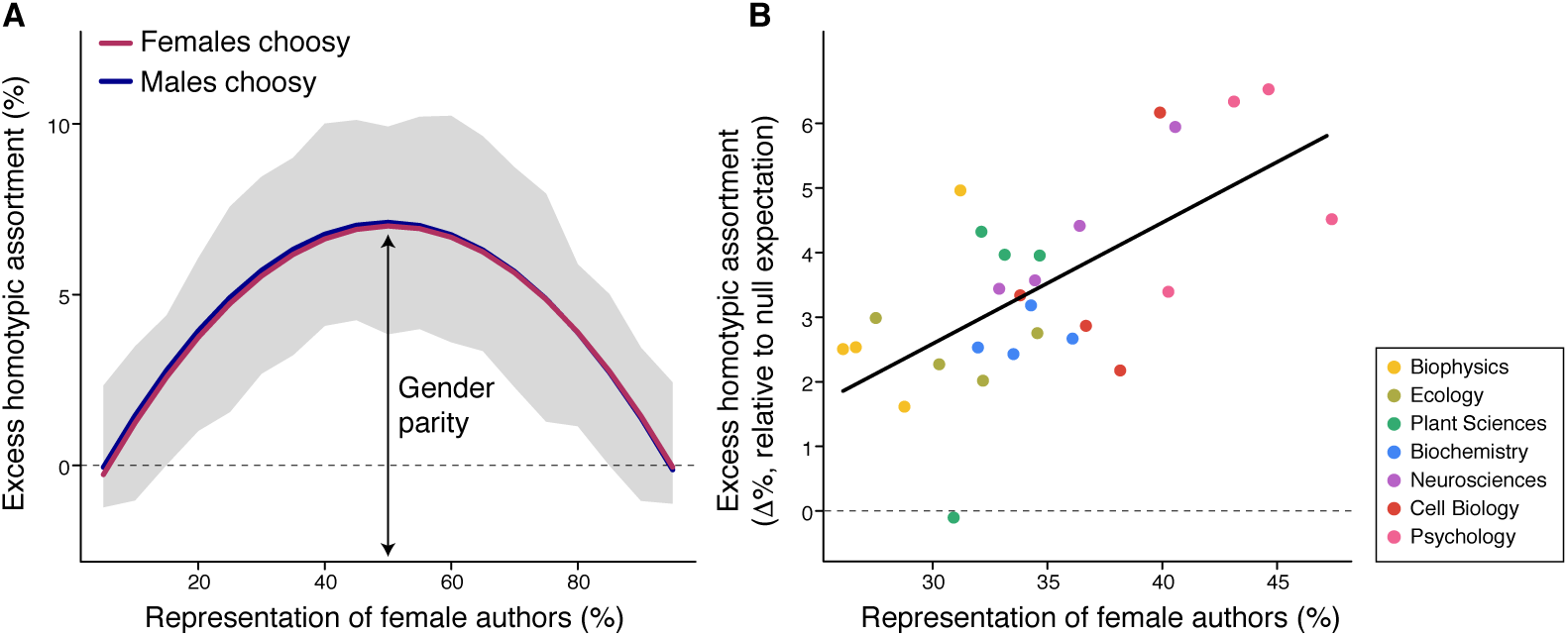
A simple model of gender homophily can explain why homotypic gender assortment increases towards parity. A) We simulated team formation when either male, or female, authors had a constant two-fold greater preference for same-gender partners (n = 3,800 simulation runs). In both cases, assortment peaked at gender parity, and then decreased according to a symmetrical polynomial relationship. The lines of best fit were determined using generalized linear regression. The shaded area is the 95% central range of excess assortment binned along the x-axis. B) Consistent with the simulation, the assortment of actual scientific authorship teams also increased towards gender parity. The scatterplot shows partial residuals (y-axis) from a mixed-effects model that accounts for variation among disciplines (Table S2).

Why should more equitable disciplines have greater homotypic gender assortment? Our simulation reveals that it is not necessarily because the individuals in more gender-balanced disciplines have stronger preferences. Instead, it is simply easier to fulfill one’s latent social preferences in a diverse environment, regardless of whether those preferences are conscious or not. Hence, the data are consistent with a scenario of widespread gender homophily that is more easily achieved in a gender-diverse field. This raises the possibility that homophily may also explain broader patterns of disciplinary representation shown in Fig. 1A. It is important to note that our bibliometric analysis is limited to a pool that is already female-deficient, because most undergraduate female trainees do not persist through training at an advanced level [1,2]. There is a strong need for additional research to evaluate whether homophily explains why many individuals drop out of science altogether [17].

Together, these results indicate that gender assortment in scientific publishing is driven by social selection during team formation, and not by gender-specific topic preferences. Based on our data, we cannot determine whether homophily is driven mainly by females who prefer to work with females, males who prefer to work with males, or both. Nor can we determine whether gender preferences in academia are exerted by first authors (typically the trainees), last authors (typically the principal investigators, PIs), or both. Recent evidence suggests that both male and female PIs may be biased towards favoring male trainees [5,18], which would be expected to counteract broader patterns of assortment. Given the small proportion of total trainees who apply to any one PI, we hypothesize that social selection by junior trainees may be the crucial factor explaining representation at both the team and disciplinary level. Remarkably, gender homotypic assortment has also been found to occur among established researchers interacting in scientific peer editing and peer reviewing networks [19], demonstrating that social selection continues to shape academia, even at advanced career stages.

To evaluate whether researchers gain an impact advantage from working in gender-matched teams, we analyzed the citation rates of >61,000 articles that had multiple authors and identifiable first- and last-author gender. We find that male gender-matched teams enjoy a significant citation advantage (Fig. 4). Females, on the other hand, are at a citation disadvantage when they work with other females (Fig. 4). The significant difference between male and female gender-matched teams corresponds to a 1.6-fold greater probability that the male-male teams will become highly cited. If we assume no gender differences in the average quality of the research, these citation disparities could be caused by bias in the research community, whereby male-led research is viewed more favorably [20]. But citation differences could also be caused by the fact that male researchers are more productive, and have greater career longevities, publication rates, and self-citation rates [1,11,21] (the latter of which could easily be due to productivity). Regardless of the mechanism, our results in Fig. 4 suggest that female researchers pay a significant citation cost to work in gender-matched teams.

**Figure 4.**
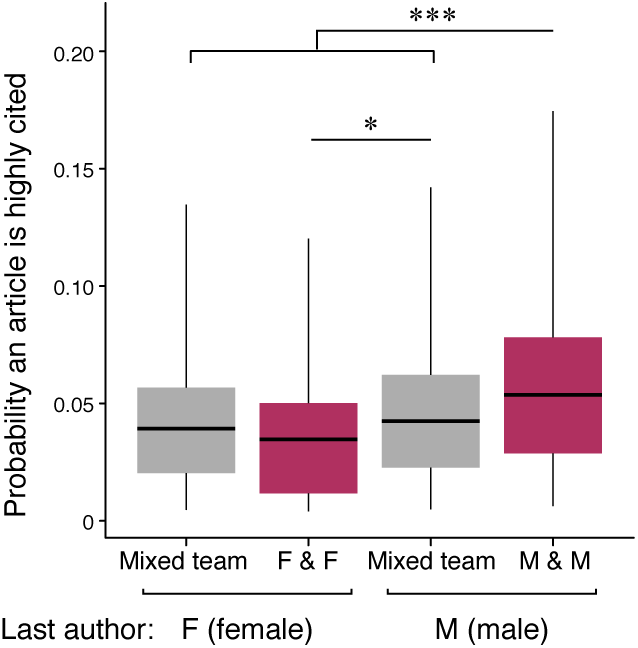
The gender composition of teams is associated with citation impact. To consider the potential effects of team composition on citation rates, we analyzed the probability that a published article became highly cited at a rate > 10 times per year (n = 61,300 articles from 7 disciplines). Male-biased teams enjoyed a significant citation advantage over other teams. In contrast, females payed a citation cost to work in gender-matched teams. Boxplots show the median, quartiles, and range of predicted probabilities (grey: mixed-gender, red: matched-gender). Asterisks indicate statistical significance in Tukey post-hoc comparisons (* p < 0.05; ** p < 0.01; *** p < 0.0001; see Table S3).

Although beyond the scope of our study, female gender-matched teams may accrue other benefits unrelated to citation rates. For example, an environment that is made up of predominantly female trainees and male senior PIs might reinforce the perception that scientific leadership is associated with maleness, regardless of the quality or intentions of the PIs [17]. Such an environment might also exacerbate stereotype threat (i.e., where females’ performance fulfills their stereotyped expectations [22]). Therefore, it is important to bear in mind that quotas or attempts to achieve gender parity through broad incentives may have unintended consequences.

Our data and simulation indicate that gender homophily is a widespread phenomenon that structures the academic environment. Given that gender differences in the attitudes towards STEM disciplines are apparent as early as kindergarten in elementary school [reviewed in 6], we propose that social preferences may amplify early disparities as students progress through educational milestones. Homophily may thus interact with other factors to ultimately produce disparities at the undergraduate level [2] and beyond (Fig. 1A). Unfortunately, women scientists are less visible to students at all levels, not only because there are fewer of them, but also because they are less widely promoted through awards and honors [23,24]. A major question is whether and how gender homophily can be harnessed to begin to have positive effects on the STEM career pipeline. This is important because our data predict that the extent of homotypic gender assortment may increase as fields become increasingly gender-balanced in the future.

## Acknowledgements

We thank Julie Morand-Ferron for insightful discussion and feedback.

## Notes

**Competing Interests:** We declare no competing interests.

## REFERENCES

1. Larivière V, Ni C, Gingras Y, Cronin B, Sugimoto CR. Bibliometrics: global gender disparities in science. Nat News. 2013;504: 211. doi:10.1038/504211a

2. American Physical Society, Integrated Postsecondary Education Data System. Bachelor’s Degrees Earned by Women, by Major [Internet]. Available: https://www.aps.org/programs/education/statistics/womenmajors.cfm

3. Eddy SL, Brownell SE, Wenderoth MP. Gender gaps in achievement and participation in multiple introductory biology classrooms. CBE—Life Sci Educ. 2014;13: 478–492. doi:10.1187/cbe.13-10-0204

4. Adamo SA. Attrition of women in the biological sciences: workload, motherhood, and other explanations revisited. BioScience. 2013;63: 43–48. doi:10.1525/bio.2013.63.1.9

5. Moss-Racusin CA, Dovidio JF, Brescoll VL, Graham MJ, Handelsman J. Science faculty’s subtle gender biases favor male students. Proc Natl Acad Sci. 2012;109: 16474–16479. doi:10.1073/pnas.1211286109

6. Ceci SJ, Ginther DK, Kahn S, Williams WM. Women in academic science: a changing landscape. Psychol Sci Public Interest. 2014;15: 75–141. doi:10.1177/1529100614541236

7. Ceci SJ, Williams WM, Barnett SM. Women’s underrepresentation in science: sociocultural and biological considerations. Psychol Bull. 2009;135: 218–261. doi:10.1037/a0014412

8. Liénard JF, Achakulvisut T, Acuna DE, David SV. Intellectual synthesis in mentorship determines success in academic careers. Nat Commun. 2018;9: 4840. doi:10.1038/s41467-018-07034-y

9. Fortunato S, Bergstrom CT, Börner K, Evans JA, Helbing D, Milojević S, et al. Science of science. Science. 2018;359: eaao0185. doi:10.1126/science.aao0185

10. West JD, Jacquet J, King MM, Correll SJ, Bergstrom CT. The role of gender in scholarly authorship. PLOS ONE. 2013;8: e66212. doi:10.1371/journal.pone.0066212

11. King MM, Bergstrom CT, Correll SJ, Jacquet J, West JD. Men set their own cites high: gender and self-citation across fields and over time. Socius. 2017;3: 2378023117738903. doi:10.1177/2378023117738903

12. Salerno PE, Páez-Vacas M, Guayasamin JM, Stynoski JL. Male principal investigators (almost) don’t publish with women in ecology and zoology. PLOS ONE. 2019;14: e0218598. doi:10.1371/journal.pone.0218598

13. McPherson M, Smith-Lovin L, Cook JM. Birds of a feather: homophily in social networks. Annu Rev Sociol. 2001;27: 415–444. doi:10.1146/annurev.soc.27.1.415

14. Kegen NV. Science networks in cutting-edge research institutions: gender homophily and embeddedness in formal and informal networks. Procedia - Soc Behav Sci. 2013;79: 62–81. doi:10.1016/j.sbspro.2013.05.057

15. Main JB. Gender homophily, Ph.D. completion, and time to degree in the humanities and humanistic social sciences. Rev High Educ. 2014;37: 349–375. doi:10.1353/rhe.2014.0019

16. Robins G, Elliott P, Pattison P. Network models for social selection processes. Soc Netw. 2001;23: 1–30. doi:10.1016/S0378-8733(01)00029-6

17. Moss-Racusin CA, Sanzari C, Caluori N, Rabasco H. Gender bias produces gender gaps in STEM engagement. Sex Roles. 2018;79: 651–670. doi:10.1007/s11199-018-0902-z

18. Sheltzer JM, Smith JC. Elite male faculty in the life sciences employ fewer women. Proc Natl Acad Sci. 2014;111: 10107–10112. doi:10.1073/pnas.1403334111

19. Helmer M, Schottdorf M, Neef A, Battaglia D. Gender bias in scholarly peer review. eLife. 2017;6: e21718. doi:10.7554/eLife.21718

20. Knobloch-Westerwick S, Glynn CJ, Huge M. The Matilda Effect in science communication: an experiment on gender bias in publication quality, perceptions and collaboration interest. Sci Commun. 2013;35: 603–625. doi:10.1177/1075547012472684

21. Zeng XHT, Duch J, Sales-Pardo M, Moreira JAG, Radicchi F, Ribeiro HV, et al. Differences in collaboration patterns across discipline, career stage, and gender. PLOS Biol. 2016;14: e1002573. doi:10.1371/journal.pbio.1002573

22. Spencer SJ, Steele CM, Quinn DM. Stereotype threat and women’s math performance. J Exp Soc Psychol. 1999;35: 4–28. doi:10.1006/jesp.1998.1373

23. Lincoln AE, Pincus S, Koster JB, Leboy PS. The Matilda Effect in science: awards and prizes in the US, 1990s and 2000s.Soc Stud Sci. 2012;42: 307–320. doi:10.1177/0306312711435830

24. Schroeder J, Dugdale HL, Radersma R, Hinsch M, Buehler DM, Saul J, et al. Fewer invited talks by women in evolutionary biology symposia. J Evol Biol. 2013;26: 2063–2069. doi:10.1111/jeb.12198

