## Supporting Information for "Gender bias in research teams and the underrepresentation of women in science"

### Article data

We used the Web of Science Core Collection database (Clarivate Analytics) to compile bibliometric data on scientific articles published in four years (2007, 2011, 2015 and 2018). All searches were performed in March and April of 2019. We started sampling articles from 2007, because that was the earliest year when full first names of authors were commonly available, facilitating analysis of gender. We chose to study biological disciplines, because the research communities within biology show consistent (stable) differences in female representation, ranging from around 25% female to gender parity (Fig. 1A). Moreover, females often make up the majority of biology students at the lowest academic levels (e.g., undergraduate students; [1]), making it an especially pertinent field for examining the causes of underrepresentation. We chose seven of the major disciplines within biology listed in Fig. 1A (corresponding to "Research Areas" in the Web of Science user interface). Each of these disciplines was chosen because it represents a research community with its own journals, meetings, and professional societies. We limited our search to articles in the Web of Science database that were associated with at least one address within the United States, England, or Canada, and we compiled up to the first 5,000 articles found for each combination of year, country, and discipline.

All data processing steps and analyses were performed in R [2]. The initial set of articles contained duplicates, because articles were often attributed to more than one of the countries and/or disciplines. If an article was found under more than one of the Research Area searches, we assigned it a single primary discipline based on the Research Area that was listed first in its Web of Science category field code ("WC"). After checking for and removing duplicate articles that had the same title and author list, the total number of unique articles was 105,558 (see Table S1), representing about 10% of the collective output for the United States, England, and Canada in the chosen disciplines and study years. We defined the primary sub-discipline of each article as its first Web of Science category entry that was distinct from its Research Area, if such data were available.

### Gender

To study the composition of research teams, we first filtered the data to the 101,524 articles that had at least two authors. We focused on the first and last authors of these articles, because by convention in biology these two authorship positions represent the largest intellectual contributors to a research article. In multi-authored articles, the first author is almost always a student or junior collaborator who led most stages of the research. The last author position is reserved for the principle investigator (PI) who conceived of and co-led the research, and also provided mentorship, funding, and infrastructure.

We used publicly-available data on US births to determine author gender probabilities based on first names [3]. The US birth name dataset represents a large sample of English name spellings for a wide range of ethnicities and years, and hence can be used to estimate gender in large-scale studies when surveys and other forms of direct and/or self-identification are not possible [4]. We used data on birth years from 1945 to 1994 (inclusive), yielding a sample of over 180.1 million first name-gender assignments. For each of the 54,543 unique first names in this dataset of 180.1 million births, we calculated the proportion of babies given that name that were male. This yielded an estimate of the probability that a given name represents a male vs. a female.

The majority (89%) of first names in the birth dataset had gender probabilities that were highly certain (i.e., all births with that name were male, or all of them were female). The remaining 11% of names were gender-uncertain to varying degrees. To deal with this uncertainty, we assigned a confidence threshold of >0.8 probability; an author's name had to be either >80% male, or >80% female, to assign the gender. In this way, we were able to determine gender for over 72% of the first and last authors of multi-author teams (147,177 first and last author assignments among 101,524 articles with more than one author). Authors whose names did not meet the 80% threshold were classified as unknown gender and omitted from further gender analyses. For each discipline-year, we used the pooled sample of first and last authors to calculate female representation in the corpus, as the percentage of all author instances assigned a gender that were female. We estimated the stability of female representation by calculating the intraclass correlation coefficient (repeatability) and its 95% confidence interval using the ICC package [5].

**Team analysis**

We used the distribution of first and last author genders within each discipline to compute null expectations for gender assortment at the collective (disciplinary) level. Because first and last author positions signify different career stages and levels of prestige in biology [4], the null computation was based on the gender ratios in each of the two authorship positions. In other words, our null model preserved seniority differences that are meaningful in biological publishing. To perform the calculation, we used the observed ratios in each discipline to compute the expected probabilities of all four team combinations (F&M, F&F, M&F, or M&M) under the assumption that coauthor selection was random. This null model was strongly rejected for 25/28 discipline-years (chi-squared tests, p ranging from < 0.0001 to 0.02 for 25 of the discipline-years). The only exceptions were Plant Sciences in 2007 ($p = 0.98$), Biophysics in 2015 ($p = 0.10$), and Cell Biology in 2018 ($p = 0.07$). Notably, each of these disciplines had author teams that were significantly more assorted than expected in all other years.

To define a measure of homotypic gender assortment on teams that would comparable across disciplines, we added the probabilities F&F + M&M for each discipline-year. This metric gives the percentage of articles that exhibited gender matching among major authors. We used a paired t-test to compare the observed and expected values of homotypic gender assortment ($n = 28$ discipline-years). To consider sources of variation across disciplines, we calculated the excess assortment by taking the observed – the expected value. We used a Gaussian mixed-effects model in the lme4 package [6] to analyze whether excess assortment could be explained by female representation; the model also included a random effect of discipline to account for other unmeasured differences. Note that because the sample size of 28 discipline-years was small, we were unable to consider many parameters in this analysis. We verified that including publication year as either a fixed or random effect significantly reduced the goodness-of-fit of the model (all $\Delta$AIC > 2).

**Network analysis**

We used network analyses to test the hypothesis that females and males may prefer different research topics within disciplines. The first step was to develop a network that would form a landscape of articles within each discipline, connected by their shared keywords. To achieve the most accurate network topology, we included both very broad and relatively specific keywords. To do this, we compiled each article's unique entries under the Web of Science field codes "DE", "ID", and "WC". These three field codes indicate the author-provided keywords, the Web of Science Keywords Plus ® (Clarivate Analytics), and Web of Science categories, respectively. The DE and ID keywords are mainly lower-level (narrower) topics of research, whereas the Web of Science category provides the sub-discipline(s) for each article. For computational feasibility during the link-building phase, we filtered any rare keywords that did not apply to at least 1% of articles within a discipline (n = 83–140 common keywords retained per discipline). Any keywords that were synonymous with the discipline itself were also excluded, because they applied to all articles in the network.

To build the networks, we constructed an unweighted and undirected adjacency matrix with each article represented on both axes. Two articles were linked with an entry of 1 if they shared at least one keyword, and 0 if they did not share any keywords. We used the igraph package [7] to create networks from these adjacency matrices, wherein each article is represented by a single node. To analyze the network topology, we used Newman's assortativity coefficient r, which is a correlation coefficient measuring the extent to which nodes with similar properties cluster together [8]. The first step was to verify that the networks had a detectable topic structure, as intended. To do this, we tested the assortment (r) of the primary sub-discipline data. Note that this step necessarily omitted articles that did not have a sub-discipline identified (range 0-51% of articles omitted per network). Given that the networks formed from the remaining articles all had significant positive sub-discipline structure ($0.2 < r < 0.7$; network sizes ranging from 4,102–12,543 articles, see Fig. 2), the next step was to examine whether the author genders were nonrandomly distributed on the network topology. These analyses were conducted for first and last author gender separately, and omitted articles where author gender was not known. Error bars were determined for all assortment estimates by bootstrapping the networks to remove 5% of the original nodes, followed by recalculating the assortment statistic, for 500 iterations.

As an additional check of our results, we also examined four annual networks formed by combining all articles from all disciplines in each year. These annual networks were also positively assorted by the discipline (all $r > 0.32$), but not by first or last author gender (all $r < 0.02$). Finally, we note that bibliometric studies often use citation networks. However, given that citations are also affected by prestige and other factors, including gender (e.g., [9,10] and Fig. 4 of this study), we deemed that networks based on keywords provided the best means to test the topic preference hypothesis. This is because citations also represent a form of prestige and influence.

**Homophily simulation**

We used an individual-based simulation model to explore what occurs if researchers have moderate preferences for same-gender teammates. In each simulation run, n = 1,000 trainees sought to be mentored by one of n = 1,000 mentors. Each mentor could accept a maximum of one trainee. Prior to the start of the simulation, genders were randomly assigned to the 2,000 individuals using a fixed ratio (selected from the range of 5–95% female, and applied to both the trainee and mentor pools). To simulate a spatially structured scenario whereby trainees are only aware of a limited set of potential mentors, we constrained each trainee to a selection of up to 40 candidate mentors. We also constrained trainees that were nearby to one another in the simulated geography to have overlapping (but never identical) sets of candidate mentors. Each trainee chose to apply to one mentor from his or her candidate set. We simulated two main scenarios: a "females choosy" scenario, wherein female trainees had a two-fold greater probability of selecting a female mentor (if available) as opposed to a male. Male trainees selected randomly in this first scenario. Alternatively, in the "males choosy" scenario, the male trainees had a two-fold greater probability of selecting a male from their candidate (if available) as opposed to a female (and female trainees chose randomly). We randomized the order in which trainees chose their mentors, removing the selected mentors from the pool with each step. We ran 100 replicate simulations at each of the 19 representation options (5%, 10%, 15% ... to 95%), for a total of n = 3,800 iterations.

**Citation analysis**

To analyze citation rates, we defined highly cited articles as those that had accrued 10 or more citations per year (on average) since the year when the article was published. The number of years since publication was calculated by subtracting the publication year from 2019. We fit a binomial mixed-effects model of the binary response variable (highly cited or not) in the lme4 package [6]. The model included random effects of discipline and publication year, as well as fixed effects of the composition of first + last authors (four levels: F&M, F&F, M&F and M&M) and publication year (standardized to have mean 0, SD 1, to facilitate the model-fitting algorithm). We used Tukey's post-hoc tests to compare all possible author composition scenarios using the multcomp package [11]. We determined that the conclusions of this analysis were robust to alternative specifications of publication year in the model syntax.

**Data availability**

All data and code necessary to reproduce our results are available at: https://figshare.com/s/c43abb9e5f755962dfd4

1. Eddy SL, Brownell SE, Wenderoth MP. Gender gaps in achievement and participation in multiple introductory biology classrooms. CBE—Life Sci Educ. 2014;13: 478–492. doi:10.1187/cbe.13-10-0204

2. R Core Team. R 3.5.1: A Language and Environment for Statistical Computing [Internet]. Vienna, Austria: R Foundation for Statistical Computing; 2018. Available: https://www.R-project.org

3. US Social Secutiry Administration. National Data [Internet]. 2015. Available: https://www.ssa.gov/oact/babynames/limits.html

4. West JD, Jacquet J, King MM, Correll SJ, Bergstrom CT. The role of gender in scholarly authorship. PLOS ONE. 2013;8: e66212. doi:10.1371/journal.pone.0066212

5. Wolak M. ICC 2.3.0: facilitating estimation of the intraclass correlation coefficient [Internet]. 2015. Available: https://cran.r-project.org/web/packages/ICC/index.html

6. Bates D, Maechler M, Bolker B, Walker S, Christensen RHB, Singmann H, et al. lme4 1.1-18-1: linear mixed-effects models using “Eigen” and S4 [Internet]. 2018. Available: https://cran.r-project.org/web/packages/lme4/index.html

7. Csardi G, coathors. igraph 1.2.2: network analysis and visualization. [Internet]. 2018. Available: https://cran.r-project.org/web/packages/igraph/index.html

8. Newman MEJ. Mixing patterns in networks. Phys Rev E. 2003;67: 026126. doi:10.1103/PhysRevE.67.026126

9. Aksnes DW, Rorstad K, Piro F, Sivertsen G. Are female researchers less cited? A large-scale study of Norwegian scientists. J Am Soc Inf Sci Technol. 2011;62: 628–636. doi:10.1002/asi.21486

10. Larivière V, Ni C, Gingras Y, Cronin B, Sugimoto CR. Bibliometrics: global gender disparities in science. Nat News. 2013;504: 211. doi:10.1038/504211a

11. Hothorn T, Bretz F, Westfall P, Heiberger RM, Schuetzenmeister A, Scheibe S. multcomp 1.4-10: simultaneous inference in general parametric models [Internet]. 2019. Available: https://cran.r-project.org/web/packages/multcomp/index.html

**Table S1. Sample sizes for each biological discipline.** Totals were summed across the four study years. Average representation was calculated as the mean representation of females in the first and last author position over the four study years.

| Discipline (WOS Research Area) | Total number of articles | Number of articles analyzed for team composition* | Size of the article network** | Average representation of female authors (%) |
|---|---|---|---|---|
| Biophysics | 9,783 | 4,667 | 4,102 | 28.2 |
| Ecology | 18,328 | 11,951 | 11,554 | 31.1 |
| Plant Sciences | 13,353 | 6,508 | 5,486 | 32.7 |
| Biochemistry Molecular Biology | 15,743 | 8,958 | 7,796 | 34.0 |
| Neurosciences | 22,267 | 13,311 | 12,543 | 36.1 |
| Cell Biology | 12,268 | 6,459 | 6,059 | 37.1 |
| Psychology | 13,816 | 9,454 | 8,900 | 43.9 |
| AVERAGE | 15,080 | 8,758 | 8,063 | 34.7 |

* To be retained in this analysis, an article had to have at least two authors with the first and last author genders identified with > 80% probability.

** To be retained in this analysis, an article had to have at least two authors with the first and last author genders identified with > 80% probability, and at least one keyword or "Web of Science Category" that was shared with >1% of the articles in its discipline. Keywords were compiled from the Web of Science field codes "DE", "ID", and "WC".

**Table S2. Analysis of excess homotypic gender assortment in relation to female author representation within a discipline (n = 28 discipline-years).** The response variable is the percent of articles that had gender-assorted author teams in excess of the null expectation. The model includes a random effect of discipline (7 levels).

| Fixed effect | Estimate (SE) | 95% CI | t-value | p-value |
|---|---|---|---|---|
| Female representation | 0.19 (0.05) | [0.08, 0.29] | 3.56 | 0.006 |

**Table S3. Analysis of the probability that an article became highly cited in relation to team composition (n = 61,300 articles).** Note that an article had to have at least two authors, each of whom had their gender identified with > 80% probability, to be included in this analysis. The model includes random effects of discipline and publication year. P-values for gender composition (a four-level factor) are reported here comparing each level with “M & F” teams. Note that in the main results, we report post-hoc p-values from a test that performed of all possible comparisons of these four factor levels.

| Fixed effect | Estimate (SE) | 95% CI | z-value | p-value |
|---|---|---|---|---|
| Year (standardized) | –0.67 (0.22) | [–1.10, –0.25] | –3.09 | 0.002 |
| Gender composition* | | | | |
| M & F | -- | -- | -- | -- |
| F & F | –0.13 (0.08) | [–0.29, 0.03] | –1.63 | 0.10 |
| F & M | 0.06 (0.07) | [–0.07, 0.19] | 0.93 | 0.35 |
| M & M | 0.31 (0.06) | [0.19, 0.43] | 4.99 | < 0.0001 |

* Stated as [first author] & [last author]
